# Investigating the Causal Role of Motor Brain Areas in Rhythm Reproduction: A Transcranial Direct Current Stimulation Study

**DOI:** 10.1101/2023.10.02.560212

**Authors:** Marina Emerick, Joshua Hoddinott, Jessica A. Grahn

## Abstract

Humans have an intrinsic tendency to move to music. However, our understanding of the neural mechanisms underlying the music-movement connection remains limited, and most studies have used correlational methods. Here, we used transcranial direct current stimulation (tDCS) to investigate the causal role of four brain regions commonly involved in movement timing and beat perception: the supplementary motor area (SMA), left and right premotor cortices (PMC), and the right cerebellum. On three different days, subjects received anodal, cathodal, or sham stimulation while reproducing strong-beat, weak-beat, and non-beat rhythms via finger tapping. Each subject only received stimulation in one of the four brain regions. As the SMA appears to play a primary role in beat perception, while the premotor cortex and cerebellum appear to have a more general role in timing, we predicted that the SMA stimulation would affect reproduction of rhythms with a beat, whereas premotor and cerebellar stimulation would affect reproduction of sequences with no beat. As expected, reproduction accuracy depended on beat strength; strong-beat rhythms were more accurately reproduced than weak and non-beat rhythms. Unexpectedly, tDCS had no effect on reproduction accuracy in any brain region. Thus, we found no evidence that modulating brain excitability in SMA, PMC, or cerebellum altered accuracy of rhythm reproduction. We discuss the implications of these results and future perspectives for this research.

## Introduction

Humans spontaneously synchronize movements to music. Although some movements, like dancing and playing a musical instrument, are complex, others, like foot tapping and nodding heads, occur spontaneously, and without training (Repp & Su, 2013). Spontaneous synchronization to music aligns with the steady pulse, or “beat” underlying the regular rhythm. While rhythm can be defined as “the serial pattern of variable note durations in a melody” (Schulkind, 1999), the feeling of a recurring pattern of salient pulses is defined as the beat (Levitin et al., 2018).

The behavoural link between rhythm and the motor system has been supported by neuroimaging studies of auditory rhythm (Bengtsson et al., 2009; Chen et al., 2008a; Grahn & Brett, 2007; Grahn & Rowe, 2009; Grahn & Schuit, 2012; Hoddinott et al., 2021; Kasdan et al., 2022; Schubotz et al., 2000). In addition to auditory sensory regions, rhythm perception activates motor areas, including the supplementary motor area (SMA), premotor cortices (PMC), the basal ganglia, and the cerebellum, even in tasks requiring no movement (Bengtsson et al., 2009; Chen et al., 2008a; Grahn & Brett, 2007; Kornysheva et al., 2010; Schubotz et al., 2000). When rhythms have a clear, or strong, beat, the basal ganglia and SMA show greater activation than when rhythms have no beat (Grahn & Brett, 2007). While the SMA and basal ganglia seem to be beat sensitive, other regions in the rhythm-perception network such as the PMC, prefrontal cortex, inferior parietal lobule, and cerebellum do not seem to be as sensitive to the beat. These regions also exhibit greater activity for complex rhythms, in which the beat is difficult to detect (Grahn & Rowe, 2009). It has been proposed that the SMA and dorsal striatum are responsible for structuring beat-based temporal anticipation and auditory expectations, creating a loop that facilitates beat perception through the entrainment of neural activity and auditory-motor interactions (Cannon & Patel, 2021; Patel & Iversen, 2014).

Neuropsychological work with Parkinson’s disease (PD) patients has highlighted the importance of the basal ganglia in beat perception, as PD patients can serve as a model for basal ganglia dysfunction. Compared to healthy controls, PD patients perform worse than controls in discriminating strong-beat rhythms, but not weak-beat rhythms, indicating the role of the basal ganglia in beat perception (Grahn & Brett, 2009). In contrast, patients with cerebellar degeneration show impairments in absolute timing, which involves encoding the durations of time intervals (McAuley & Jones, 2003; Teki et al., 2011), but perform similarly to healthy controls in beat-based timing tasks (Breska & Ivry, 2018). Other studies support that the cerebellum appears to respond and encode temporal intervals that do not have a reference interval, such as a regular beat (Nozaradan et al., 2017; Povel & Essens, 1985; Teki et al., 2011). Overall, these studies support a dissociation between the basal ganglia in beat-based, or relative, timing and the cerebellum in absolute timing.

Despite extensive research using correlational neuroimaging methods (Grahn & Brett, 2007; Grahn & Rowe, 2009; Teki et al., 2011), and neuropsychological studies (Breska & Ivry, 2018; Grahn & Brett, 2009), few studies have used causal methods in healthy humans (Leow et al., 2022). Transcranial direct current stimulation (tDCS) is one way to causally examine the role of different brain areas in different timing processes. TDCS modulates synaptic efficacy of neurons by altering resting membrane potential by passing a weak electric current between two brain areas (Purpura & McMurtry, 1965), and it can modulate brain responses in two directions: anodal stimulation increases cortical excitability, while cathodal stimulation inhibits cortical excitability (Reinhart et al., 2017; Thair et al., 2017). While tDCS does not directly trigger action potentials, it brings neural populations closer to or further from activation threshold, increasing or decreasing the likelihood of neural activity, respectively. Importantly, tDCS is functionally specific – behavioural changes can be seen if a modulated network is task-relevant (Bikson & Rahman, 2013).

A previous study using strong- and weak-beat rhythms demonstrated the SMA’s crucial role in rhythm discrimination using tDCS (Leow et al., 2022). During anodal stimulation of the SMA, participants discriminated rhythm changes more accurately than during sham stimulation, and during cathodal stimulation, participants discriminated more poorly than during sham. This polarity-dependent response is strong evidence for role of the SMA in rhythm discrimination, however, the effect was observed for both strong- and weak-beat rhythms. Premotor cortex stimulation elicited no consistent effect on discrimination performance. For the cerebellum, both anodal and cathodal stimulation worsened discrimination performance. These results indicate that both the SMA and cerebellum play roles in rhythm discrimination, but, they do not support a selective role of the SMA role in beat-based timing, as stimulation affected discrimination of both strong- and weak-beat rhythms (Leow et al., 2022). Further, the cerebellar stimulation had a selective effect on strong-beat rhythm discrimination, counter to its proposed role in absolute, not relative (beat-based) timing.

Here, we follow up the rhythm discrimination results with a different measure of beat perception. We used a rhythm reproduction task for multiple reasons. First, it can often provide a more sensitive and continuous measure of rhythm and beat perception accuracy, as well as minimizing decisional effects (e.g., response bias) and serial position effects of the to-be-discriminated change in the rhythm present in perception tasks (Grahn & Brett, 2007; Grahn & Rowe, 2009; Leow et al., 2022). Second, the PMC group did not show effects of tDCS in the prior rhythm discrimination study (Leow et al., 2022), but its possible PMC plays a role when programming rhythmic motor output. Finally, we wanted to test whether SMA stimulation would also affect performance when a motor output action was required (Cannon & Patel, 2021; Patel & Iversen, 2014).

We also included non-beat rhythms along with strong- and weak-beat rhythms used previously. This enabled us to determine whether the lack of difference between strong- and weak-beat rhythms for the SMA stimulation group would replicate with non-beat rhythms (Leow et al., 2022). In non-beat rhythms, no beat can be consistently aligned to the rhythm, as the intervals are irregular. In weak-beat rhythms, a beat theoretically can be aligned to the rhythm (although, in practice, performance of weak- and non-beat rhythms is similar, unless they are presented in a looping fashion). Moreover, its possible that the cerebellar contributions will be larger for non-beat (more irregular) rhythms compared to weak-beat rhythms.

Here, we investigated the causal role of four brain areas (SMA, right cerebellum, left PMC, and right PMC) in rhythm and beat perception through a rhythm reproduction paradigm, assessing how reproduction accuracy differed for sequences that could be timed using a beat-based versus non-beat-based timing system when neural excitability was causally altered using tDCS. A mixed design was employed because of significant individual differences in rhythm reproduction ability (Grahn & Schuit, 2012) and tDCS responsivity (Chew et al., 2015). Participants were randomly assigned to one of four brain area groups and completed the rhythm reproduction task on three different days while receiving sham, anodal, or cathodal tDCS stimulation.

Based on neuroimaging evidence showing the SMA is involved in the temporal processing of beat-based sequences and that the premotor cortex and cerebellum appear to respond in both beat and non-beat contexts (Grahn & Brett, 2007) or respond more to non-beat-based contexts (Breska & Ivry, 2018; Teki et al., 2012), we hypothesize that the SMA plays a primary role in beat perception. Thus, modulating excitability in the SMA should influence the ability to reproduce strong-beat rhythms more than other rhythm types. In contrast, if the cerebellum and PMC are involved in absolute timing, effects of tDCS will not depend on beat strength, and their stimulation should influence accuracy for all rhythm types, or more so for weak and non-beat rhythms than strong-beat rhythms.

Findings from this study will shed light on rhythm and beat perception by identifying the causal role of specific brain regions in beat-based and non-beat based timing systems during rhythm reproduction. It will clarify whether the SMA is selectively part of the beat-based timing system, and whether the cerebellum and PMC are part of the non-beat-based timing system. We are building on neuropsychological studies by stimulating areas that are not commonly dysfunctional in clinical populations, such as the SMA and PMC, using healthy controls.

## Methods

### Participants

Participants were primarily recruited through the Western University undergraduate participant pool (SONA) or through word of mouth. The Health Sciences Research Ethics Board at Western University approved the study, and the experiments were performed following relevant guidelines and regulations.

To minimize potential risks, participants were excluded if they had a history of psychiatric or neurological problems such as epileptic seizures, Tourette’s syndrome, ADHD, depression; any metallic implants, such as pacemakers, cerebral aneurysm clips or other electronic implants; any active skin problems, such as eczema; any unstable medical condition and the susceptibility to migraine or other frequent headaches; any history of episodes of faintness; current use of a hearing aid; for female participants specifically, being pregnant, or trying to become pregnant. All participants gave their informed consent before beginning the experiment.

In total, 91 participants took part in the study. Ten participants were excluded for different reasons, such as not completing all three sessions, feeling uncomfortable in the active session, or technical issues. Therefore, the final sample consisted of 81 participants (age mean ± standard deviation: 19.7 ± 3.9, age range: 18-41, 54 women); 22 participants in the SMA stimulation group, 19 in the right cerebellar stimulation group, 19 in the left PMC stimulation group, and 21 in the right PMC stimulation group. Participants were additionally categorized into groups of high musical experience and low musical experience depending on their scores from the Goldsmith MSI musical training subscale (Müllensiefen et al., 2014). Because the scores’ median split was not exactly, we arbitrarily divided participants with ranges from 21 to 49 as high musical experienced participants, and ranges from 7 to 20 as low musical experienced participants. This yielded to 42 participants with high musical experience (ranges from 21 to 44) and 39 participants with low musical experience (ranges from 7 to 20), however, they were not evenly split across groups: there were fourteen high musical experience participants in the SMA group, ten in the cerebellum group, eight in the left PMC group, and ten in the right PMC group.

## Material

### Questionnaires

Musical training ability was assessed with the musical training subscale from the Goldsmiths Musical Sophistication Index (Müllensiefen et al., 2014), a self-report questionnaire that evaluates musical sophistication as a multidimensional construct. The subscale comprises seven items, and its score can range from 7 to 49. A demographic questionnaire with questions about educational level, language, and general health was used.

A questionnaire was developed following Schaal et al. (2021) to assess participants’ awareness of the type of stimulation. By the end of each session, participants were asked to indicate whether they thought they had received active or sham stimulation. If they indicated active, they indicated whether they thought it was anodal or cathodal stimulation. They also indicated how sure they were on an adapted Likert scale, with 1 = ‘completely unsure’ and 10 = ‘completely sure’. Finally, they were also asked if they noticed any sensation difference (e.g., tingling, itching sensation) during or after the stimulation. Participants were not aware that sham was a possibility during the experiment, but they were free to answer the after-session questionnaire how they wanted to. If they did not know the meaning of any term, they were told that they would receive explanation about it by the end of the study.

### Stimuli

Stimuli were presented using E-prime 2.0 Software (Psychology Software Tools, Pittsburgh, PA) on a Dell laptop. Participants listened to the auditory stimuli through Bose headphones. Rhythms were generated using Matlab Software (Matlab, 2016).

Stimuli comprised rhythms adapted from the previous work of Grahn & Brett (2007). Rhythms were separated into three categories according to their beat strength: strong-beat, weak-beat, and non-beat rhythms. The ‘perceptual accents’, or the feeling that a note is more prominent than its surrounding notes, causing the beat to be salient, was manipulated in each type of rhythm (Povel & Essens, 1985; Povel & Okkerman, 1981). While strong- and weak-beat rhythms consist of integer-ratio intervals (e.g., 1:2:3:4), non-beat rhythms consist of non-integer ratios where the ‘2’ and ‘3’ intervals are replaced by ‘1.4’ and ‘3.6’ respectively (i.e., 1:1.4:3.6:4). Strong-beat rhythms induce regularly occurring perceptual accents at the beginning of each group of four units (e.g., ‘1111’, ‘22’, or ‘4’), emphasizing the beat at predictable intervals. Weak-beat rhythms have more irregular accents, and less tones occurring on the beat, making the beat less emphasized and hard to detect. In addition to irregularly spaced accents, in non-beat rhythms the intervals themselves are irregular, making a consistent beat impossible (see Figure 1).

**Figure 1.**
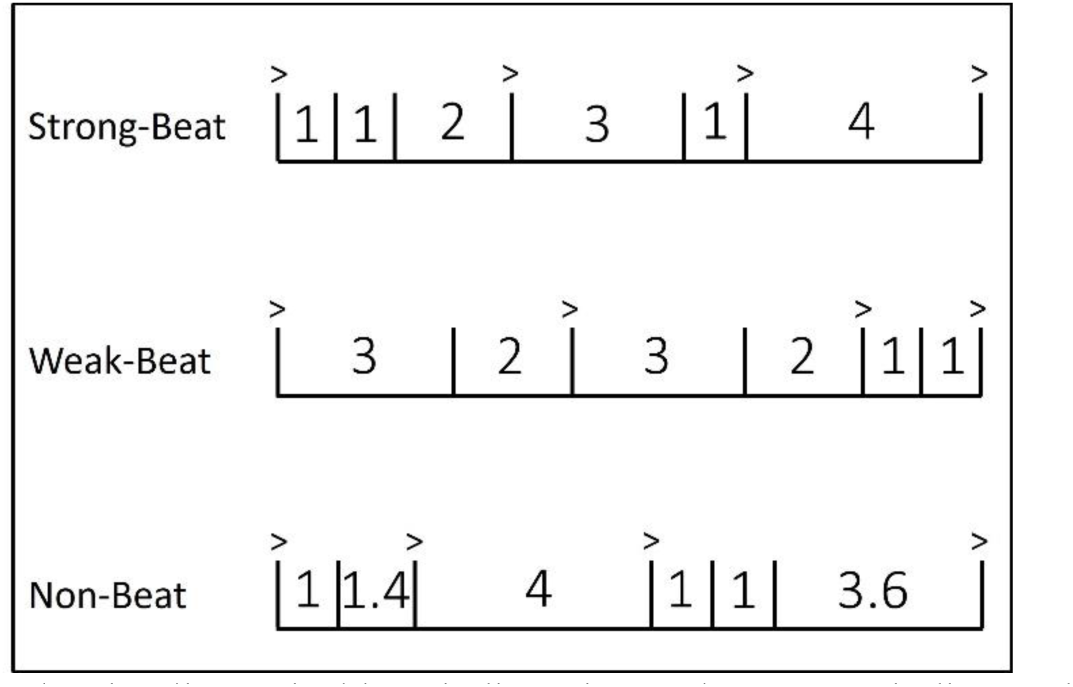
Schematic of sample stimuli. Vertical bars indicate interval onset, ‘>’ indicate where perceptual accents should be heard (Povel & Okkerman, 1981). Numbers indicate the relationship between intervals. The base interval, or the shortest interval (e.g. ‘1’) could go from 225 ms to 275 ms, in steps of 25 ms. The other intervals are multiples of the base interval, for example, ‘2’ is twice the duration of ‘1’ (Adapted from Grahn and Brett 2007, and Hoddinott & Grahn, in Prep).

Each rhythm comprised 5 to 7 intervals, and interval durations were multiples of the ‘base interval’, or shortest interval (i.e., ‘1’ interval). Base intervals could be either 225, 250 or 275 ms, in order to eliminate any carryover effects of a perceived beat rate from one trial to another. For example, for a strong-beat rhythm with a base interval of 250 ms, the sequence 112314 contains the intervals ‘250 250 500 750 250 1000’ in length (milliseconds). To ensure that the length of each reproduced interval in the rhythm could be measured, each rhythm ended with an additional tone equal to the ‘1’ interval that marked the end of the final interval. For example, a six-interval rhythm would have seven tone onsets, and thus seven tap times to determine the duration of those six intervals. Additionally, for each beat type, the stimuli comprised six rhythms with five intervals, seven rhythms with six intervals and seven rhythms with seven intervals. The set of intervals used to create each rhythm was termed an ‘interval set’, and the same interval set (e.g., the interval set 11334) appeared across the three rhythm types the same number of times. In total, we included 60 rhythms: 20 per rhythm type (see Table 2 for the complete list of rhythms).

**Table 1.**
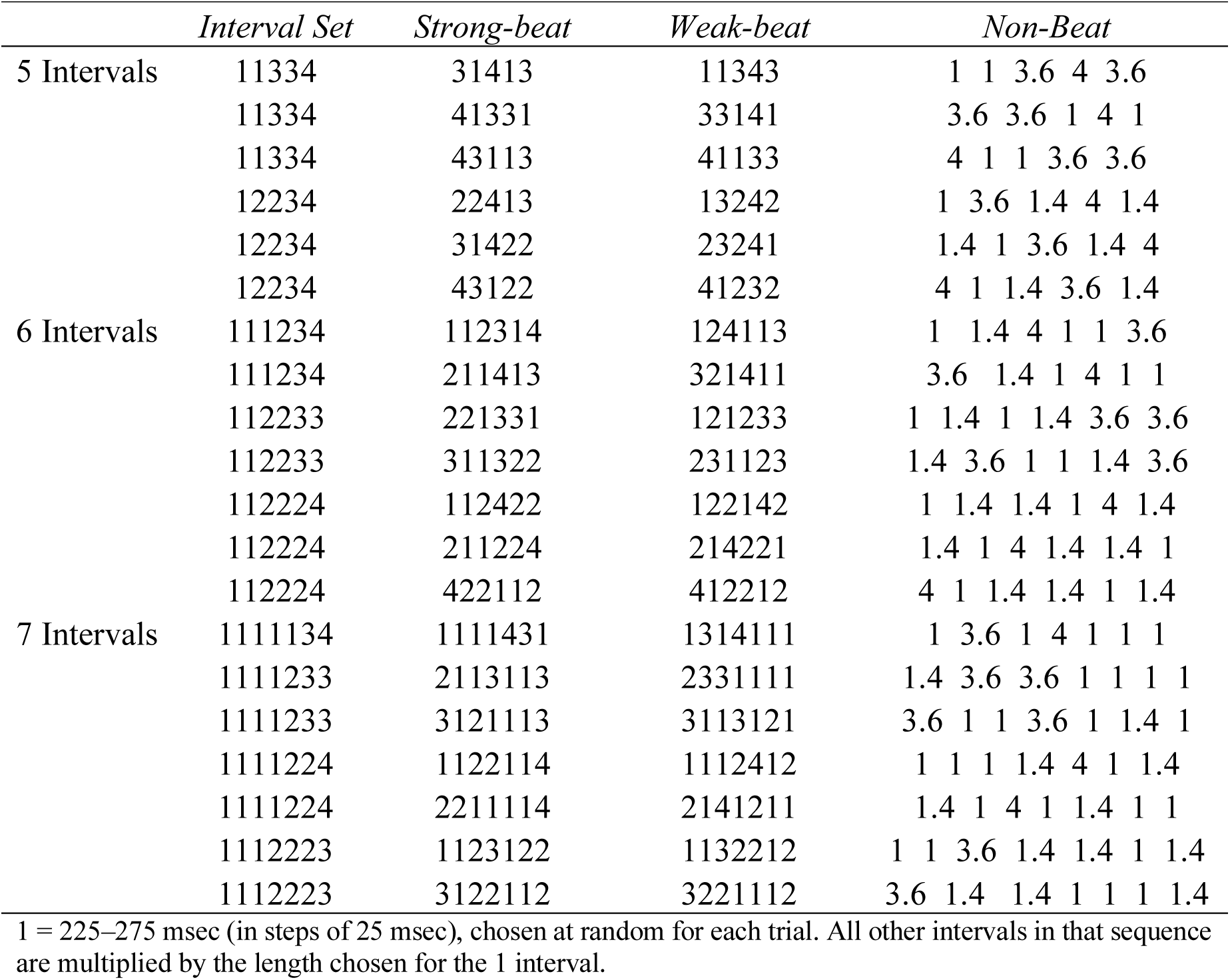
Rhythmic Sequences for Each Condition.

|  | *Interval Set* | *Strong-beat* | *Weak-beat* | *Non-Beat* |
| --- | --- | --- | --- | --- |
| 5 Intervals | 11334 | 31413 | 11343 | 1 1 3.6 4 3.6 |
|  | 11334 | 41331 | 33141 | 3.6 3.6 1 4 1 |
|  | 11334 | 43113 | 41133 | 4 1 1 3.6 3.6 |
|  | 12234 | 22413 | 13242 | 1 3.6 1.4 4 1.4 |
|  | 12234 | 31422 | 23241 | 1.4 1 3.6 1.4 4 |
|  | 12234 | 43122 | 41232 | 4 1 1.4 3.6 1.4 |
| 6 Intervals | 111234 | 112314 | 124113 | 1 1.4 4 1 1 3.6 |
|  | 111234 | 211413 | 321411 | 3.6 1.4 1 4 1 1 |
|  | 112233 | 221331 | 121233 | 1 1.4 1 1.4 3.6 3.6 |
|  | 112233 | 311322 | 231123 | 1.4 3.6 1 1 1.4 3.6 |
|  | 112224 | 112422 | 122142 | 1 1.4 1.4 1 4 1.4 |
|  | 112224 | 211224 | 214221 | 1.4 1 4 1.4 1.4 1 |
|  | 112224 | 422112 | 412212 | 4 1 1.4 1.4 1 1.4 |
| 7 Intervals | 1111134 | 1111431 | 1314111 | 1 3.6 1 4 1 1 1 |
|  | 1111233 | 2113113 | 2331111 | 1.4 3.6 3.6 1 1 1 1 |
|  | 1111233 | 3121113 | 3113121 | 3.6 1 1 3.6 1 1.4 1 |
|  | 1111224 | 1122114 | 1112412 | 1 1 1 1.4 4 1 1.4 |
|  | 1111224 | 2211114 | 2141211 | 1.4 1 4 1 1.4 1 1 |
|  | 1112223 | 1123122 | 1132212 | 1 1 3.6 1.4 1.4 1 1.4 |
|  | 1112223 | 3122112 | 3221112 | 3.6 1.4 1.4 1 1 1 1.4 |

1 = 225–275 msec (in steps of 25 msec), chosen at random for each trial. All other intervals in that sequence are multiplied by the length chosen for the 1 interval.

### Tasks

Rhythm Reproduction Task: While the stimulation/sham was applied for 20 minutes, rhythms were presented in random order for each participant. Participants passively listened to the same rhythm three times, and then reproduced what they heard by tapping it back with their index finger on a computer keyboard. Immediately after their response, a new rhythm was presented, leading to 60 trials: 20 per rhythm type. Before each session, 4 training trials were included. Training rhythms were different from the ones included in the main trials, and the training was accomplished in the same way; after hearing the rhythm three times, participants reproduced it by taping it back with a computer keyboard. Self-Paced Tapping Task: In order to control for tDCS effects on motor variability, participants completed the self-paced tapping task at the beginning of each session. This task comprised a spontaneous tapping rate activity where participants were asked to tap ten times in a row at a comfortable rate, while the stimulation/sham was applied. They were told that there was no ‘right or wrong’ for this part of the task.

### Transcranial Direct Current Stimulation (tDCS)

The Chattanooga Ionto Dual Channel Electrophoresis System was used to apply a 2 mA current over the participant’s scalp with the tDCS. Two 4 x 6 cm rubber electrodes placed in saline-soaked sponges (current density of 0.083 mA/cm^2^ in each electrode; 0.9% NaCl) were secured to the scalp with rubber head straps. For the active tDCS conditions, the current was gradually ramped up to 2 mA over 30 s upon commencing the task. The stimulation remained on during the task for 20 minutes, and it ramped down at the end of the session. For the sham tDCS conditions, the stimulation was similarly ramped up over 30 s to 2 mA but then immediately ramped back down to 0 over the next 30s. The sham condition mimics the tingling or itching feeling that some participants experience when stimulation is applied. This method is sufficient to achieve blinding in stimulation-naive participants, as it evokes the sensation of being stimulated but does not lead to a neurophysiological change (Ambrus et al., 2012). Anodal and cathodal stimulation were differentiated by whether the anode or cathode electrode was placed over the region of interest. During both anodal and cathodal stimulation, the current remained at 2 mA for the duration of the task.

The stimulation sites were located using the international electroencephalographic 10-20 system (Woods et al., 2016) and following Leow et al. procedures (2022). As shown in Figure 2, for the SMA site, one electrode was positioned 2 cm anterior to Cz (located at the center of the top of the head), and the other electrode was placed on the forehead above the right eye (Vollmann et al., 2013); for the cerebellum, one electrode was positioned 3 cm right of the inion (at the back of the head), while the other one was positioned on the right buccinator muscle (facial muscle underlying the cheek) (Galea et al., 2009), for the left PMC, one electrode was positioned 2 cm rostral to C3, while the other electrode was positioned on the contralateral orbit, and finally for the right PMC, one electrode was positioned 2 cm rostral to C4, while the other electrode was positioned on the contralateral orbit (Boros et al., 2008; Nitsche et al., 2003; Picard & Strick, 2001).

**Figure 2.**
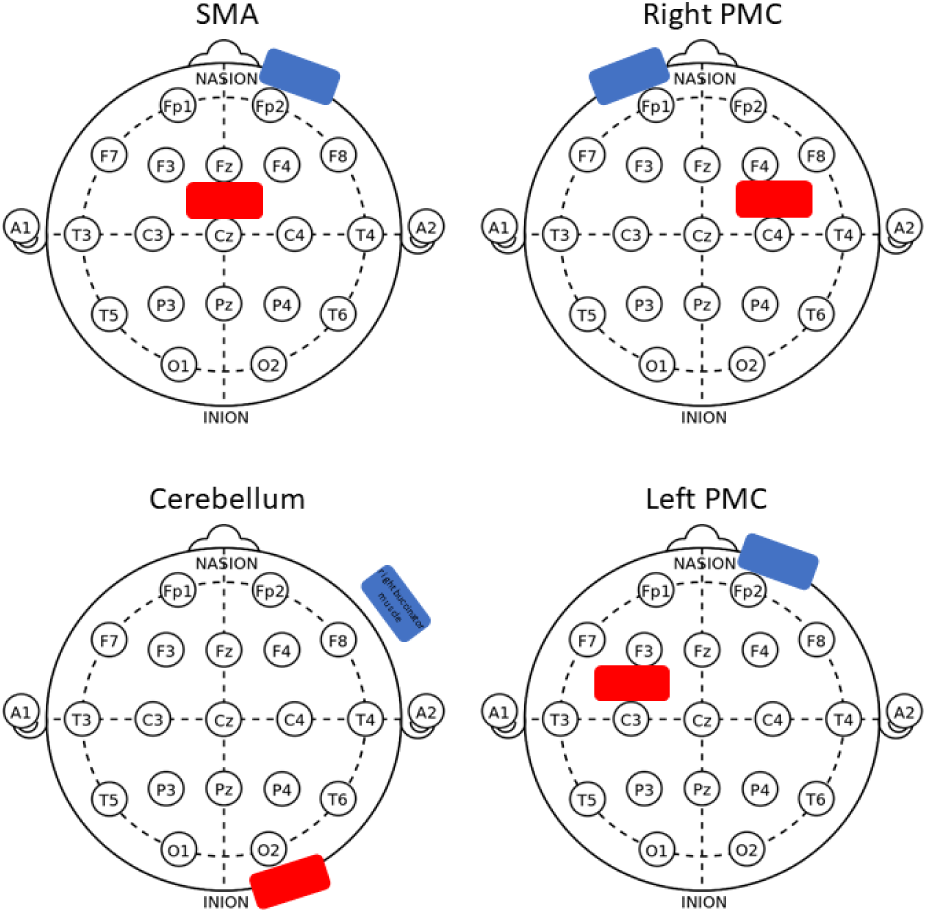
tDCS Stimulation Sites. Stimulation electrode positions, anodal electrodes in red and cathodal electrodes in blue. An anodal stimulation example is shown, as the anodal electrodes are positioned in the regions of interest, and the cathodal electrodes are positioned in the reference regions. **SMA:** anode electrode 2 cm anterior to Cz, reference above right eye forehead. **PMC:** anode electrode 2 cm anterior to C3 for right PMC, 2 cm anterior to C4 for left PMC, references in contralateral orbit. **Cerebellum:** anode 3 cm right of the inion, reference in right buccinator muscle.

## Procedure

Participants participated in three, one-hour sessions at the University of Western Ontario, with two to seven days between sessions. The three sessions were similar regarding the task procedure. The type of stimulation received (sham, anodal or cathodal) was counterbalanced across sessions, and in the first session, participants completed medical screening and demographic questionnaires as well as the musical training subscale of the Goldsmiths Musical Sophistication Index.

The self-paced tapping task was completed at the beginning of each session as a control task to assess whether the tDCS stimulation interfered with tapping responses. Afterward, participants completed the rhythm reproduction task: four practice trials followed by the main trials, where they listened to one rhythm three times and reproduced it by tapping a computer key. They reproduced each of the rhythms presented while receiving a sham or active stimulation, depending on the session. Participants completed 60 trials, reproducing 20 strong-beat rhythms, 20 weak-beat rhythms, and 20 non-beat rhythms, presented in random order. The total reproduction task lasted 20 minutes.

At the end of each session, participants were asked whether they were aware of the type of stimulation they received (active or sham). If they did not know what sham or active stimulation meant, they were told to report any different sensation they felt, and that they would receive explanation about it by the end of the study. At the end of the third day, participants were debriefed and had the opportunity to ask the experimenter any questions.

## Data Analysis

Data was first treated using R (R Core Team, 2020), and statistical analysis was performed using JASP (JASP Team, 2022).

### Rhythm Reproduction Task

#### Proportion of Correct Trials

Any trials with the incorrect number of taps (either too few or too many) were deemed incorrect and not further analyzed. For the remaining trials, the intertap time of each interval in a rhythmic sequence was compared to its corresponding presented interval in that rhythm, then a measure of the proportion of correctly reproduced trials was derived from the rhythm reproduction data. Trials with the correct number of taps (e.g., for a 6-interval rhythm, seven taps should be counted) and in which all interval durations were reproduced within 20% of the presented interval (e.g., for a 250 ms interval, a reproduced interval between 200 ms – 300 ms was accepted) were counted as a correct trial. For each participant, the proportion of correct trials was calculated for each beat type (strong-, weak- and non-beat), and for each stimulation condition (sham, anodal and cathodal). Higher values represent a better performance in the task, as lower values represent the opposite.

##### Frequentist approach

A mixed-measures ANOVA was conducted on the proportion of correct trials to investigate differences based on beat strength (strong-beat vs. weak-beat vs. non-beat rhythms) and stimulation type (sham vs. anodal vs. cathodal). Brain area stimulated (SMA, right cerebellum, right PMC or left PMC) and the musical experience (high vs. low) were included as the between-subject factors. Follow-up ANOVAs were run separately for each stimulation site to investigate differences based on beat strength (strong-beat vs. weak-beat vs. non-beat rhythms) and stimulation type (sham vs. anodal vs. cathodal) while the musical experience (high vs. low) was included as the between-subject factor. Pairwise tests were used to compare significant results.

##### Bayesian Statistics

Bayesian statistics were used to confirm the evidence for the null hypothesis. Contrary to the frequentist approach, the Bayesian approach can quantify hypotheses instead of just rejecting or accepting them. Here we used a Bayesian repeated measures ANOVA to select the best fitting model that explains our data. In the ANOVA case, Bayesian statistics consider all combinations and interactions of factors to try to explain the data behavior and the best model to fit the data. The best model represents a higher BFM, or the change in likelihood of selecting a model prior to posterior data inclusion. The larger the BF, the more support for the model fitting the data we have. Besides that, it is necessary to include the likelihood of each effect in the best model. From prior and posterior probabilities of including an effect across all models, BFincl shows which factors is more likely to explain the data, with higher BFincl showing stronger evidence for the hypothesis being tested.

#### Proportional Average Error

Additionally, another type of analysis was used to evaluate the performance on the rhythm reproduction task. The average proportional error is the mean error of all the intervals in a rhythm. The error is calculated as the absolute difference between the reproduced interval (e.g. 250 ms) and the original interval (e.g. 300 ms), divided by the original interval (|250 – 300| / 300; error = 0.17). The bigger the average error, the worse the participant’s performance.

A mixed-measures ANOVA was conducted on the proportion of average error to investigate differences based on beat strength (strong-beat vs. weak-beat vs. non-beat rhythms) and stimulation type (sham vs. anodal vs. cathodal). Brain area stimulated (SMA, right cerebellum, right PMC or left PMC) and the musical experience (high vs. low) were included as the between-subject factors.

### Self-paced tapping task

For the self-paced tapping task, the intertap time marked each interval in the tapping sequence. The standard deviation of the intervals in each sequence was taken, and intervals that fell outside two standard deviations of the mean were removed from the analysis. Then, the coefficient of variation was calculated to measure the timing variability across each tap interval. A mixed-measures ANOVA was conducted on the coefficient of variation comparing the three stimulation sessions: sham, anodal and cathodal, while the brain area stimulated (SMA, right cerebellum, right PMC or left PMC) was included as the between-subject factor.

## Results

### Rhythm Reproduction Task

Mauchly’s test of sphericity indicated that the assumption of sphericity was violated for the following factors: beat strength and for the interaction between stimulation and beat strength (*p* < .05). Due to that, we used the Greenhouse-Geisser sphericity correction values.

The four brain stimulation groups did not differ in performance (*F*(3,73) = 0.76, *p* = 0.52, *ηp2* = 0.03). However, a main effect of music experience was observed (*F*(1,73) = 10.22, *p* < 0.01, *ηp2* = 0.12), as high musical experience participants performed better than low musical experience participants (*Mdiff* = 11.29, *SE* = 3.53).

When including the brain area groups as a factor, a significant main effect of beat strength was found (*F*(1.60, 117.12) = 118.54, *p* < .001, *ηp2* = .62). Post-hoc comparisons showed that strong-beat rhythms had a higher percentage of correct trials than weak-(*Mdiff* = 23.75, *SE* = 1.95, *t* = 12.19, *p* < .001) and non-beat rhythms (*Mdiff* = 27.73, *SE* = 1.95, *t* = 14.24, *p* < .001), and weak-beat rhythms had a significantly higher percent of correct trials than non-beat rhythms (*Mdiff* = 3.98, *SE* = 1.95, *t* = 2.04, *p* = 0.04). Additionally, an interaction between beat strength and music experience was observed (*F*(1.60, 117.12) = 9.41, *p* < .001, *ηp2* = .11). Participants with high musical experience had significantly better reproduction performance for strong-beat rhythms than participants with low musical experience (*Mdiff* = 20.70, *SE* = 4.19, *t* = 4.94, p < .001), but no difference for weak-beat rhythms (*Mdiff* = 8.83, *SE* = 4.19, *t* = 2.11, *p* = .17) nor for non-beat rhythms (*Mdiff* = 4.34, *SE* = 4.19, *t* = 1.04, *p* = .90).

Independently of the brain area, there was no main effect of stimulation type (*F*(2, 146) = .77, *p* = 0.46, *ηp2* = .01). No interaction between the stimulation type and beat strength was observed (*F*(2.97, 217.14) = 0.82, *p* = 0.48, *ηp2* = .01), nor between the stimulation type and brain area being stimulated (*F*(6, 146) = 1.29, *p* = 0.26, *ηp2* = .05) nor between the stimulation type and music experience (*F*(2, 146) = 0.14, *p* = 0.87, *ηp2* = .002).

In order to look deeper into each group’s performance and make sure that no other factor impacted in the null results of the interaction between brain area and stimulation, we used independent ANOVAs for each stimulation siteFor the SMA, as shown in Figure 3, there was a main effect of beat strength (*F*(1.76, 35.12) = 14.82, *p* < .001, *ηp2* = 0.47). Strong beat rhythms had a higher percentage of correct trials than weak (*Mdiff* = 15.70, *SE* = 4.15, *t* = 3.78, *p* = .001) and non-beat rhythms (*Mdiff* = 21.95, *SE* = 4.15, *t* = 5.28, *p* < .001), but the difference between weak and non-beat rhythms did not reach significance (*Mdiff* = 6.25, *SE* = 4.15, *t* = 1.50, *p* = .14). However, no effect of stimulation on performance was observed (*F*(1.56, 122.50) = .001, *p* = .99, *ηp2* = 6.58e^-5^).

**Figure 3.**
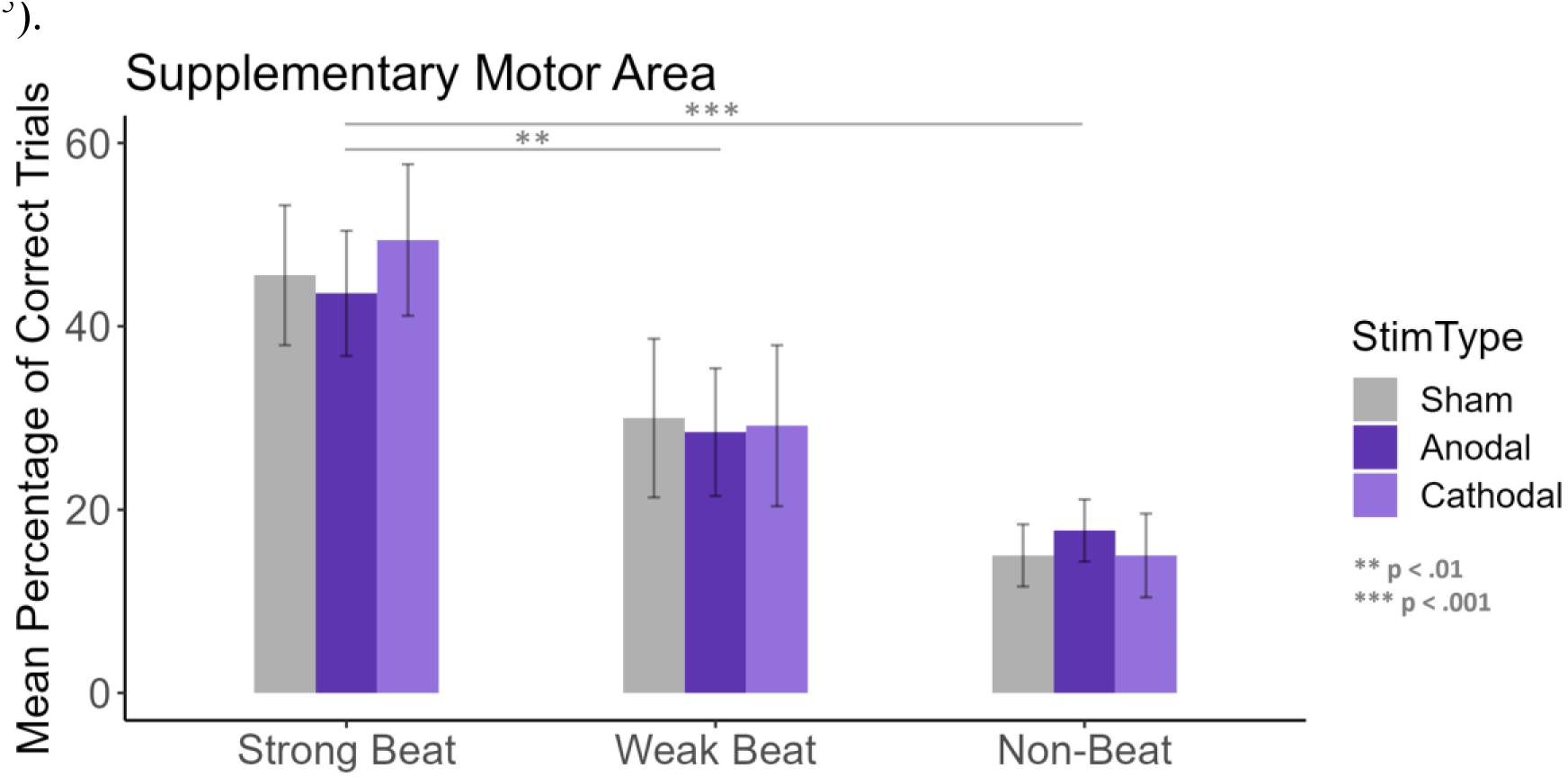
Response accuracy for the SMA group, separated according to beat strength. Stimulation type is differentiated through the different colors and hues, sham is represented in grey, anodal in darker purple and cathodal in light purple. Error bars represent the standard error of the mean. There are significant differences in accuracy when comparing beat strength but not when comparing different types of stimulation.

When separating participants according to their musical experience (nHigh = 14, nLow = 8 participants), a main effect of musical experience was observed for the SMA group (*F*(1,20) = 5.96, *p* = 0.02, *ηp2* = 0.23). A significant interaction between musical experience and beat strength was seen (*F*(1.76, 35.12) = 6.65, *p* = 0.005, *ηp2* = 0.25), with a significantly higher percentage of correct trials for high musical experience participants only for the strong-beat rhythms (*Mdiff* = 37.32, *SE* = 9.99, *t* = 3.74, *p* = .008). No interaction between musical experience and stimulation type was seen (*F*(1.57, 31.31) = 1.39, *p* = 0.26, *ηp2* = 0.06).

For the right cerebellum, as shown in Figure 4, there was a significant main effect of beat strength (*F*(1.12, 19.02) = 57.46, *p* < .001, *ηp2* = .77). Strong-beat rhythms had a significantly higher percentage of correct trials than weak-(*Mdiff* = 31.50, *SE* = 3.63, *t* = 8.68, *p* < .001) and non-beat rhythms (*Mdiff* = 35.54, *SE* = 3.63, *t* = 9.79, *p* < .001), but weak-beat rhythms did not differ from non-beat rhythms (*Mdiff* = 4.04, *SE* = 3.63, *t* = 1.11, *p* = .27). However, no effect of stimulation on performance was observed (*F*(1.87, 31.82) = 1.53, *p* = .23, *ηp2* = .08).

**Figure 4.**
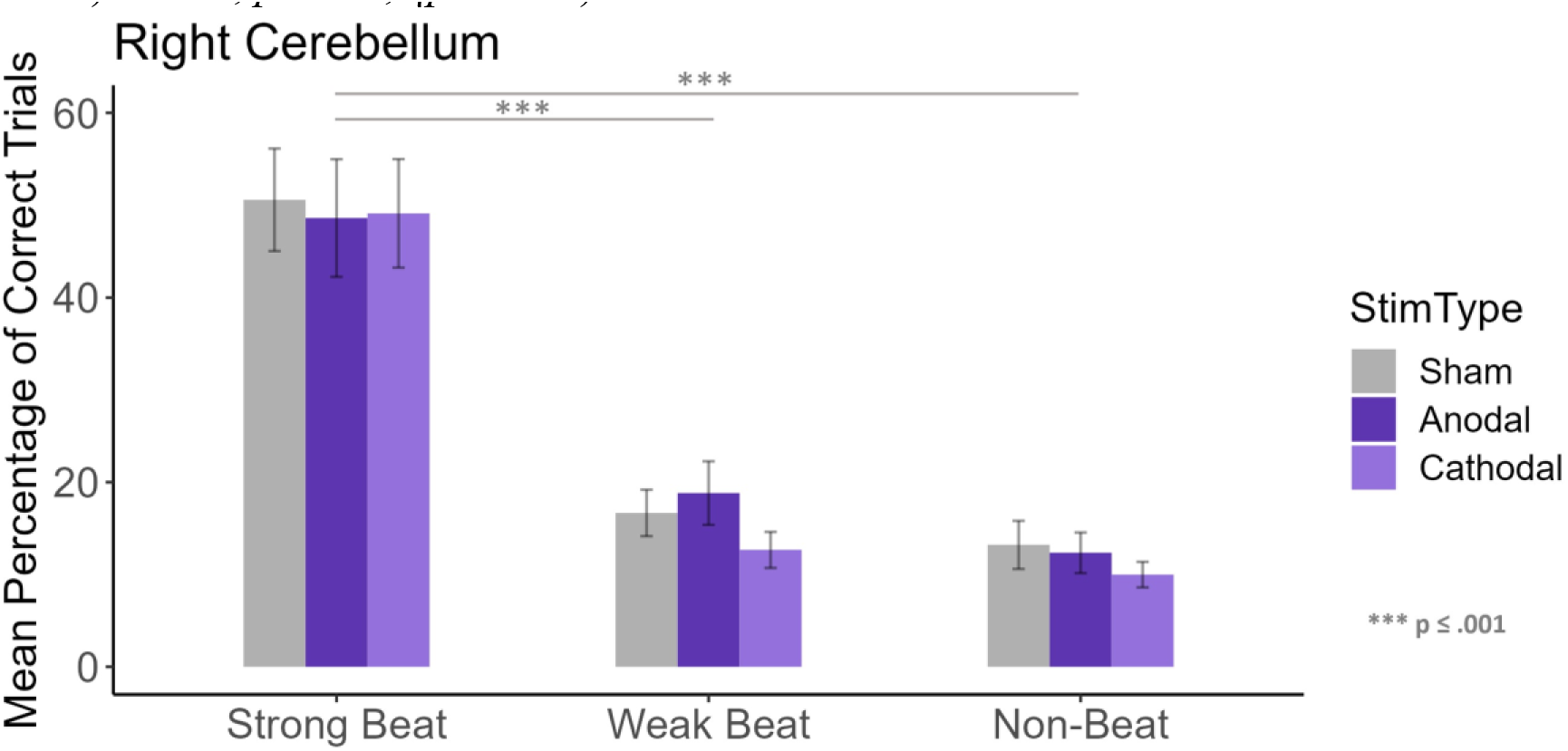
Response accuracy for the right cerebellum group, separated according to beat strength. Stimulation type is differentiated through the different purple hues, sham is represented in grey, anodal in darker purple and cathodal in light purple. Error bars represent the standard error of the mean. There are significant differences in accuracy when comparing beat strength but not when comparing different types of stimulation.

When separating participants according to their musical experience (nHigh = 10, nLow = 9 participants), there was no effect of musical experience for the right cerebellum group (*F*(1,17) = 0.78, *p* = 0.39, *ηp2* = 0.04). No interaction between musical experience and beat strength was seen (*F*(1.12, 19.02) = 0.91, *p* = 0.36, *ηp2* = 0.05), nor between musical experience and stimulation type (*F*(1.87, 31.82) = 0.42, *p* = 0.65, *ηp2* = 0.02).

For the left premotor cortex, as shown in Figure 5, there was a significant main effect of beat strength (*F*(1.16, 19.79) = 34.33, *p* < .001, *ηp2* = 0.67). Strong-beat rhythms had a significantly higher percent of correct trials than weak (*Mdiff* = 25.03, *SE* = 3.52, *t* = 7.11, *p* < .001) and non-beat rhythms (*Mdiff* = 25.50, *SE* = 3.52, *t* = 7.24, *p* < .001), but weak-beat rhythms did not differ from non-beat rhythms (*Mdiff* = 0.47, *SE* = 3.52, *t* = 0.13, *p* = 0.89). However, no effect of stimulation on performance was observed (*F*(1.97, 33.47) = 1.80, *p* = 0.18, *ηp2* = 0.09).

**Figure 5.**
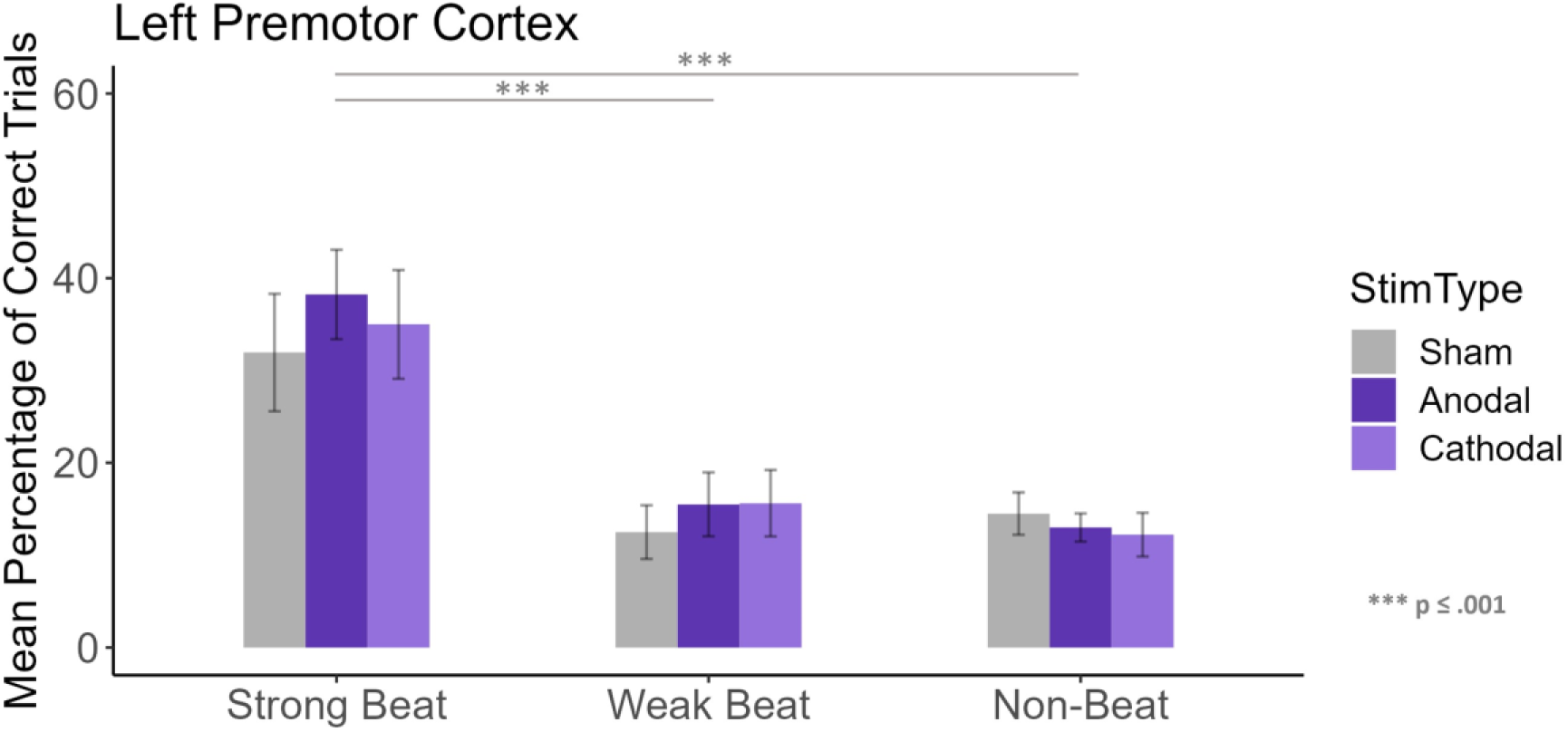
Response accuracy for the left PMC group, separated according to beat strength. Stimulation type is differentiated through the different purple hues, sham is represented in grey, anodal in darker purple and cathodal in light purple. Error bars represent the standard error of the mean. There are significant differences in accuracy when comparing beat strength but not when comparing different types of stimulation.

When separating participants according to their musical experience (nHigh = 8, nLow = 11 participants), there was no effect of musical experience for the left PMC group (F(1,17) = 1.02, p = 0.33, ηp2 = 0.06). No interaction between musical experience and beat strength was seen (F(1.16, 19.79) = 1.69, p = 0.21, ηp2 = 0.09), nor between musical experience and stimulation type (F(1.97, 33.47) = 1.99, p = 0.15, ηp2 = 0.10).

For the right premotor cortex, as shown in Figure 6, there was a significant main effect of beat strength (*F*(1.74, 33.13) = 26.72, *p* < .001, *ηp2* = 0.58). Strong-beat rhythms had a higher percent of correct trials than weak (*Mdiff* = 22.77, *SE* = 4.07, *t* = 5.60, *p* < .001) and non-beat rhythms (*Mdiff* = 27.98, *SE* = 4.07, *t* = 6.87, *p* < .001), but weak-beat rhythms did not differ from non-beat rhythms (*Mdiff* = 5.17, *SE* = 4.07, *t* = 1.27, *p* = 0.21). However, no effect of stimulation on performance was observed (*F*(1.99, 37.83) = 1.09, *p* = 0.35, *ηp2* = 0.05).

**Figure 6.**
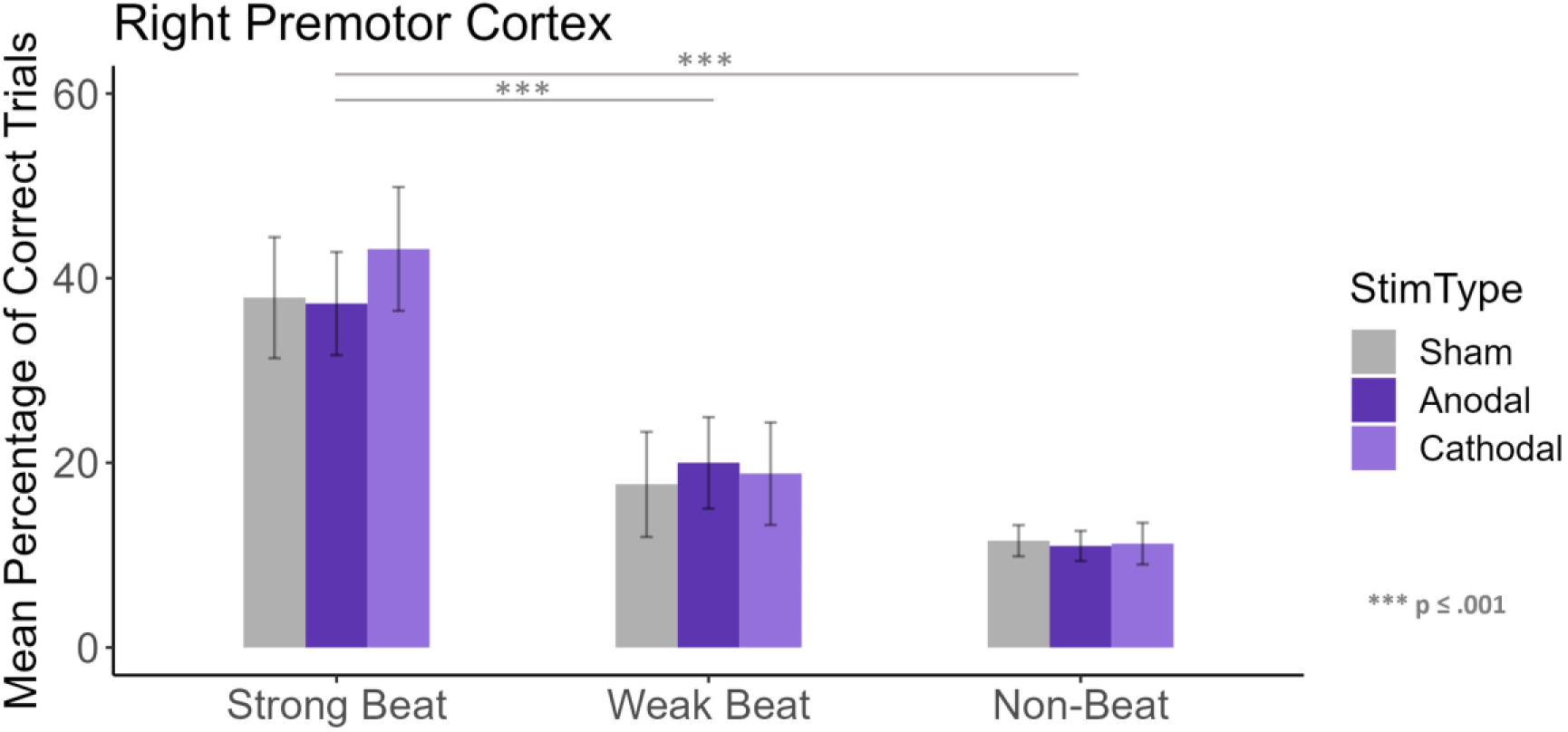
Response accuracy for the right PMC group, separated according to beat strength. Stimulation type is differentiated through the different purple hues, sham is represented in grey, anodal in darker purple and cathodal in light purple. Error bars represent the standard error of the mean. There are significant differences in accuracy when comparing beat strength but not when comparing different types of stimulation.

When separating participants according to their musical experience (nHigh = 10, nLow = 11 participants), there was no effect of musical experience for the right PMC group (*F*(1,19) = 3.52, *p* = 0.08, *ηp2* = 0.16). No interaction between musical experience and beat strength was seen (*F*(1.74, 33.13) = 2.23, *p* = 0.13, *ηp2* = 0.10), nor between musical experience and stimulation type (*F*(1.99, 37.83) = 0.22, *p* = 0.80, *ηp2* = 0.01).

### Bayesian Statistics

The best-fitting model for our data included the combination of beat strength (Beat) + brain region (Group) + musical experience (Music) + interaction of beat strength (Beat) with the brain region (Group) + interaction of beat strength (Beat) with musical experience (Music), with *BFM* = 190.56.

When analyzing the effects and including all the factors (Table 2), beat strength is very likely to explain our data (*BFincl* = 2.67e^92^), followed by the interaction between beat strength and musical experience (*BFincl* = 1.14e^9^), and by the interaction between beat strength and the brain region group (*BFincl* = 58.07). Musical experience is also likely to explain the data (*BFincl* = 20.54).

**Table 2.**
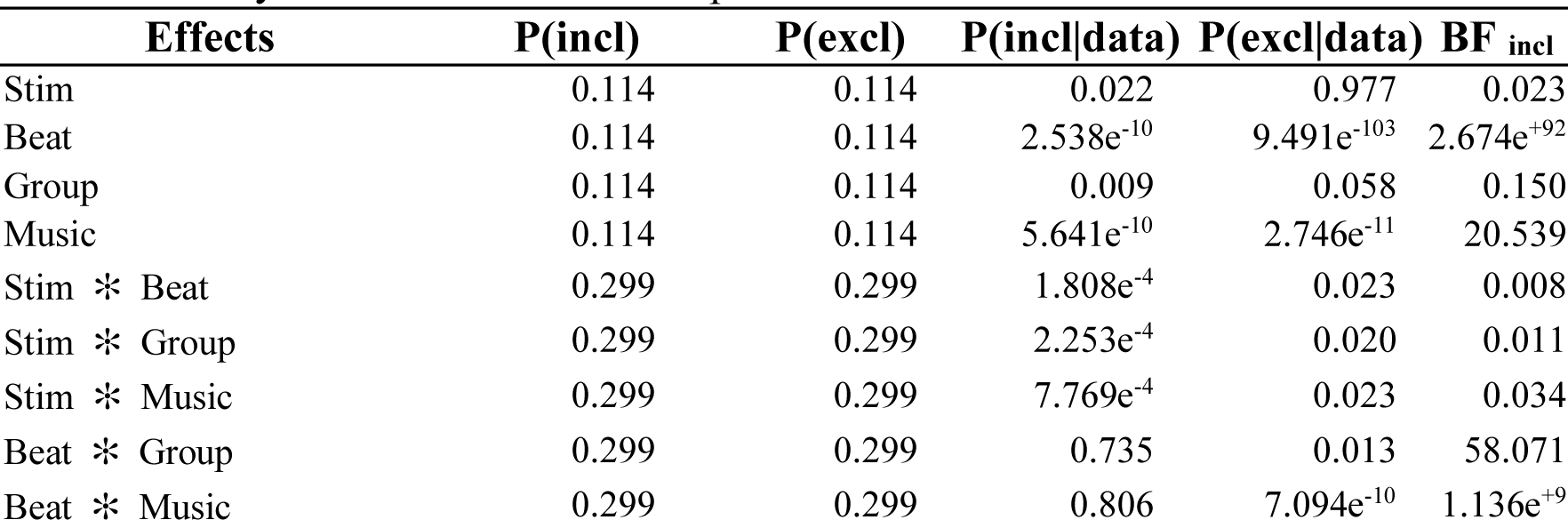
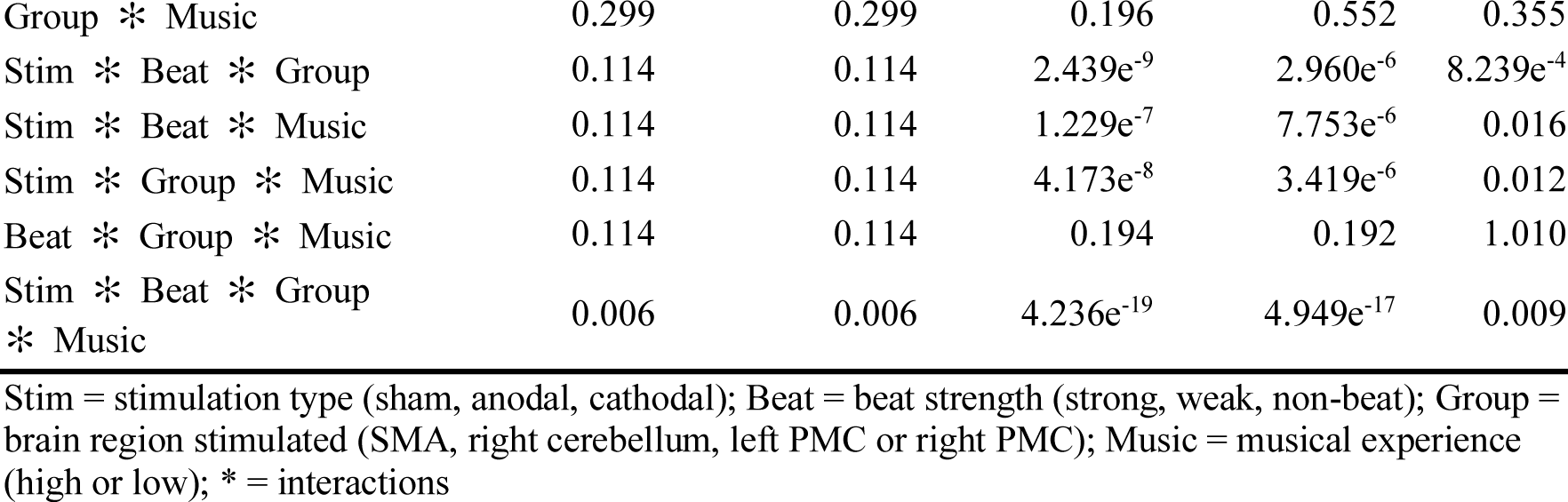
Analysis of Effects for the Proportion of Correct Trials.

| Effects | P(incl) | P(excl) | P(incl\|data) | P(excl\|data) | BF<sub>incl</sub> |
| --- | --- | --- | --- | --- | --- |
| Stim | 0.114 | 0.114 | 0.022 | 0.977 | 0.023 |
| Beat | 0.114 | 0.114 | $2.538e^{-10}$ | $9.491e^{-103}$ | $2.674e^{+92}$ |
| Group | 0.114 | 0.114 | 0.009 | 0.058 | 0.150 |
| Music | 0.114 | 0.114 | $5.641e^{-10}$ | $2.746e^{-11}$ | 20.539 |
| Stim * Beat | 0.299 | 0.299 | $1.808e^{-4}$ | 0.023 | 0.008 |
| Stim * Group | 0.299 | 0.299 | $2.253e^{-4}$ | 0.020 | 0.011 |
| Stim * Music | 0.299 | 0.299 | $7.769e^{-4}$ | 0.023 | 0.034 |
| Beat * Group | 0.299 | 0.299 | 0.735 | 0.013 | 58.071 |
| Beat * Music | 0.299 | 0.299 | 0.806 | $7.094e^{-10}$ | $1.136e^{+9}$ |
| Group * Music | 0.299 | 0.299 | 0.196 | 0.552 | 0.355 |
| Stim * Beat * Group | 0.114 | 0.114 | 2.439e<sup>-9</sup> | 2.960e<sup>-6</sup> | 8.239e<sup>-4</sup> |
| Stim * Beat * Music | 0.114 | 0.114 | 1.229e<sup>-7</sup> | 7.753e<sup>-6</sup> | 0.016 |
| Stim * Group * Music | 0.114 | 0.114 | 4.173e<sup>-8</sup> | 3.419e<sup>-6</sup> | 0.012 |
| Beat * Group * Music | 0.114 | 0.114 | 0.194 | 0.192 | 1.010 |
| Stim * Beat * Group * Music | 0.006 | 0.006 | 4.236e<sup>-19</sup> | 4.949e<sup>-17</sup> | 0.009 |

Stim = stimulation type (sham, anodal, cathodal); Beat = beat strength (strong, weak, non-beat); Group = brain region stimulated (SMA, right cerebellum, left PMC or right PMC); Music = musical experience (high or low); \* = interactions

Following the separate ANOVAs for each group, again, beat strength is the factor most likely to explain the data; for the SMA group, beat strength *BFincl* is 1.399e^14^ (Table 3), for the right cerebellum group *BFincl* is 2.431e^30^ (Table 4), for the left PMC group *BFincl* is 3.568e^23^ (Table 5), while for the right PMC group *BFincl* is 1.399e^14^ (Table 6).

**Table 3.** Analysis of effects for the proportion of correct trials - SMA group.

| Effects | P(incl) | P(excl) | P(incl\|data) | P(excl\|data) | BF<sub>incl</sub> |
| --- | --- | --- | --- | --- | --- |
| Stim | 0.737 | 0.263 | 0.071 | 0.929 | 0.027 |
| Beat | 0.737 | 0.263 | 1.000 | 2.554e<sup>-15</sup> | 1.399e<sup>+14</sup> |
| Music | 0.737 | 0.263 | 1.000 | 2.065e<sup>-7</sup> | 1.730e<sup>+6</sup> |
| Stim * Beat | 0.316 | 0.684 | 0.004 | 0.996 | 0.008 |
| Stim * Music | 0.316 | 0.684 | 0.011 | 0.989 | 0.023 |
| Beat * Music | 0.316 | 0.684 | 1.000 | 8.056e<sup>-7</sup> | 2.689e<sup>+6</sup> |
| Stim * Beat * Music | 0.053 | 0.947 | 6.966e<sup>-5</sup> | 1.000 | 0.001 |

Stim = stimulation type (sham, anodal, cathodal); Beat = beat strength (strong, weak, non-beat); Group = brain region stimulated (SMA, right cerebellum, left PMC or right PMC); Music = musical experience (high or low); \* = interactions

**Table 4.** Analysis of effects for the proportion of correct trials - right cerebellum group.

| Effects | P(incl) | P(excl) | P(incl\|data) | P(excl\|data) | BF<sub>incl</sub> |
| --- | --- | --- | --- | --- | --- |
| Stim | 0.263 | 0.263 | 0.175 | 0.804 | 0.218 |
| Beat | 0.263 | 0.263 | 0.836 | 3.438e<sup>-31</sup> | 2.431e<sup>+30</sup> |
| Music | 0.263 | 0.263 | 0.285 | 0.551 | 0.519 |
| Stim * Beat | 0.263 | 0.263 | 0.010 | 0.186 | 0.051 |
| Stim * Music | 0.263 | 0.263 | 0.012 | 0.082 | 0.144 |
| Beat * Music | 0.263 | 0.263 | 0.156 | 0.293 | 0.533 |
| Stim * Beat * Music | 0.053 | 0.053 | 3.602e<sup>-5</sup> | 1.956e<sup>-4</sup> | 0.184 |

Stim = stimulation type (sham, anodal, cathodal); Beat = beat strength (strong, weak, non-beat); Group = brain region stimulated (SMA, right cerebellum, left PMC or right PMC); Music = musical experience (high or low); \* = interactions

**Table 5.** Analysis of effects for the proportion of correct trials - left PMC group.

| Effects | P(incl) | P(excl) | P(incl\|data) | P(excl\|data) | BF<sub>incl</sub> |
| --- | --- | --- | --- | --- | --- |
| Stim | 0.263 | 0.263 | 0.097 | 0.879 | 0.110 |

**Table 4.** Analysis of effects for the proportion of correct trials - right cerebellum group.

| Effects | P(incl) | P(excl) | P(incl\|data) | P(excl\|data) | BF<sub>incl</sub> |
| --- | --- | --- | --- | --- | --- |
| Beat | 0.263 | 0.263 | 0.355 | 9.949e<sup>-25</sup> | 3.568e<sup>+23</sup> |
| Music | 0.263 | 0.263 | 0.136 | 0.218 | 0.627 |
| Stim * Beat | 0.263 | 0.263 | 0.007 | 0.114 | 0.063 |
| Stim * Music | 0.263 | 0.263 | 0.018 | 0.080 | 0.227 |
| Beat * Music | 0.263 | 0.263 | 0.643 | 0.139 | 4.611 |
| Stim * Beat * Music | 0.053 | 0.053 | 1.358e<sup>-4</sup> | 8.291e<sup>-4</sup> | 0.164 |

Stim = stimulation type (sham, anodal, cathodal); Beat = beat strength (strong, weak, non-beat); Group = brain region stimulated (SMA, right cerebellum, left PMC or right PMC); Music = musical experience (high or low); \* = interactions

**Table 6.** Analysis of effects for the proportion of correct trials - right PMC group.

| Effects | P(incl) | P(excl) | P(incl\|data) | P(excl\|data) | BF<sub>incl</sub> |
| --- | --- | --- | --- | --- | --- |
| Stim | 0.737 | 0.263 | 0.100 | 0.900 | 0.040 |
| Beat | 0.737 | 0.263 | 1.000 | 2.554e<sup>-15</sup> | 1.399e<sup>+14</sup> |
| Music | 0.737 | 0.263 | 0.955 | 0.045 | 7.654 |
| Stim * Beat | 0.316 | 0.684 | 0.004 | 0.996 | 0.009 |
| Stim * Music | 0.316 | 0.684 | 0.009 | 0.991 | 0.021 |
| Beat * Music | 0.316 | 0.684 | 0.892 | 0.108 | 17.850 |
| Stim * Beat * Music | 0.053 | 0.947 | 3.362e<sup>-5</sup> | 1.000 | 6.052e<sup>-4</sup> |

Stim = stimulation type (sham, anodal, cathodal); Beat = beat strength (strong, weak, non-beat); Group = brain region stimulated (SMA, right cerebellum, left PMC or right PMC); Music = musical experience (high or low); \* = interactions

### Proportional Average Error

Mauchly’s test of sphericity indicated that the assumption of sphericity was violated for the following factors: beat strength and for the interaction between stimulation and beat strength (*p* < .05). Due to that, we used the Greenhouse-Geisser sphericity correction values.

The four brain stimulation groups did not differ in performance (*F*(3,73) = 2.47, *p* = .07, *ηp2* = 0.09). However, a main effect of music experience was observed (*F*(1,73) = 4.38, *p* = .04, *ηp2* = .06), as high musical experience participants performed better than low musical experience participants (*Mdiff* = 5.70, *SE* = 2.72). See figure 8.

**Figure 8.**
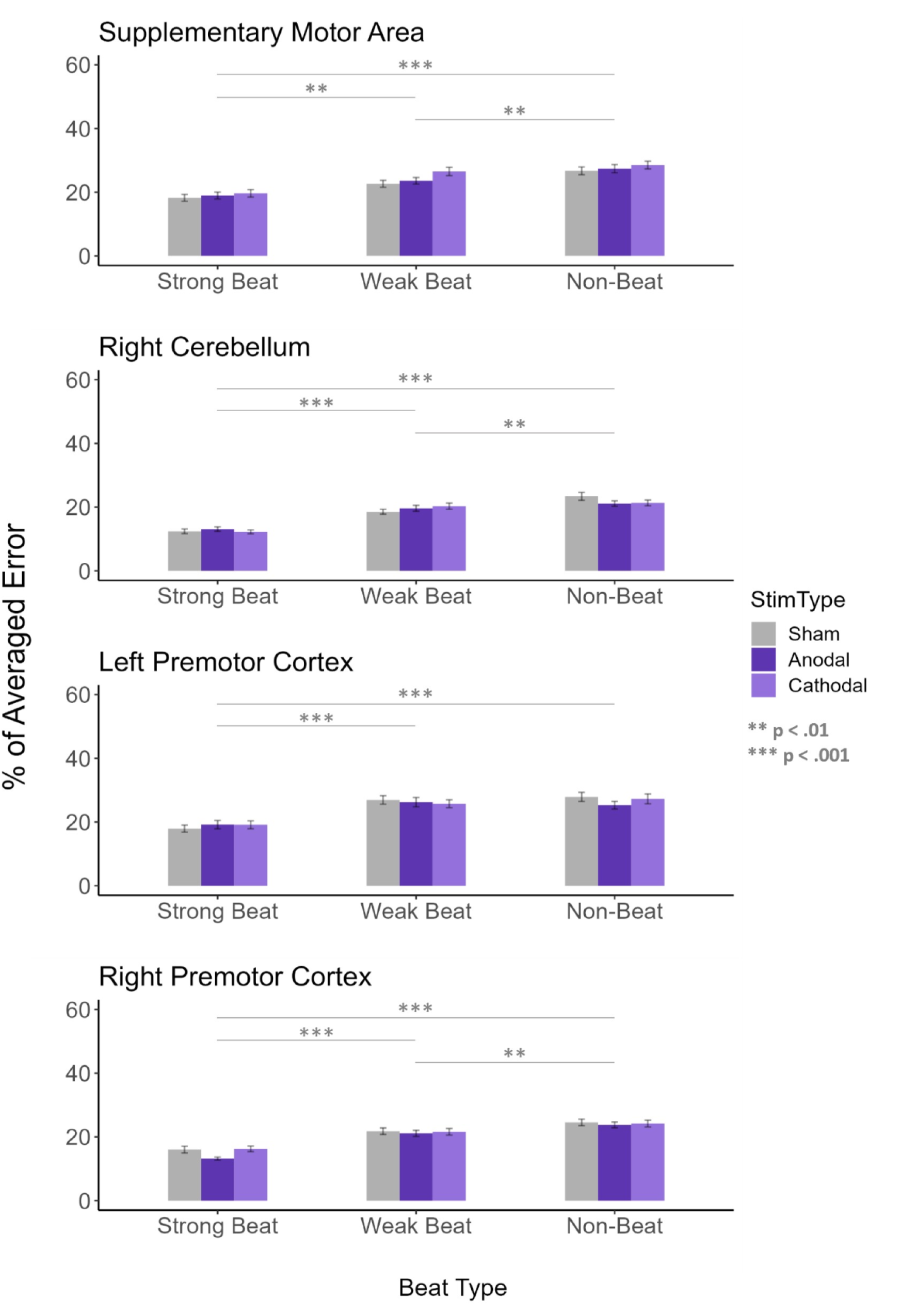
Proportion for the averaged error for the four groups (SMA, right cerebellum, left PMC and right PMC). Perfect performance would result in zero percentage of averaged error, the bigger the values, the more averaged error is shown. Average error is separated according to beat strength. Stimulation type is differentiated through the different colors and hues, sham is represented in grey, anodal in darker purple and cathodal in light purple. Error bars represent the standard error of the mean. There are significant differences in the averaged error when comparing beat strength but not when comparing different types of stimulation.

When including the brain area groups as a factor, a significant main effect of beat strength was found (*F*(1.76, 128.48) = 137.60, *p* < .001, *ηp2* = 0.65). Post-hoc comparisons showed that strong-beat rhythms had a higher percentage of correct trials than weak-(*Mdiff* = 6.62, *SE* = 0.57, *t* = 11.67, *p* < .001) and non-beat rhythms (*Mdiff* = 9.11, *SE* = 0.57, *t* = 16.05, *p* < .001), and weak-beat rhythms had a significantly higher percent of correct trials than non-beat rhythms (*Mdiff* = 2.49, *SE* = 0.57, *t* = 4.38, *p* < .001).

Independently of the brain area, there was no main effect of stimulation type (*F*(2, 146) = 0.75, *p* = 0.48, *ηp2* = 0.01). No interaction between the stimulation type and beat strength was observed (*F*(2.96, 216.01) = 0.31, *p* = .81, *ηp2* = .004), nor between the stimulation type and brain area being stimulated (*F*(6, 146) = 0.45, *p* = 0.84, *ηp2* = .02) nor between the stimulation type and music experience (*F*(2,146) = 1.36, *p* = .26, *ηp2* = .02).

### Self-Paced Tapping Task

The four brain stimulation groups did not differ in variability of self-paced tapping (F(3,73) = 0.34, p = 0.79, ηp2 = 0.01). There was no main effect of stimulation when comparing sham vs. anodal vs. cathodal sessions (F(2,146) = 0.82, p = 0.44, ηp2 = 0.01). No interaction between stimulation type and musical experience was seen (F(2,146) = 0.74, p = 0.48, ηp2 = 0.01), nor between stimulation type and brain area being stimulated (F(6,146) = 1.47, p = 0.19, ηp2 = 0.06).

### Blindness Towards Stimulation

Participants correctly guessed they received sham stimulation in 32.10% of sessions, while for anodal stimulation they correctly guessed it was an active anodal stimulation in 41.97% of sessions, and for cathodal, they correctly guessed it was an active cathodal stimulation in 23.46% of sessions. On a scale of how sure they were about their answer, with 1 = ‘completely unsure’ and 10 = ‘completely sure’, the average was 2.98 for all participants in the three sessions. The guessed percentages are above chance for the anodal and sham sessions, but they could only indicate that participants were guessing with uncertainty and luckily answering it right given their low average of certainty in the certainty scale. Besides that, 49.38% of the sham sessions were incorrectly guessed as active sessions. These results indicate that participants were probably blind towards stimulation type.

**Figure 5.**
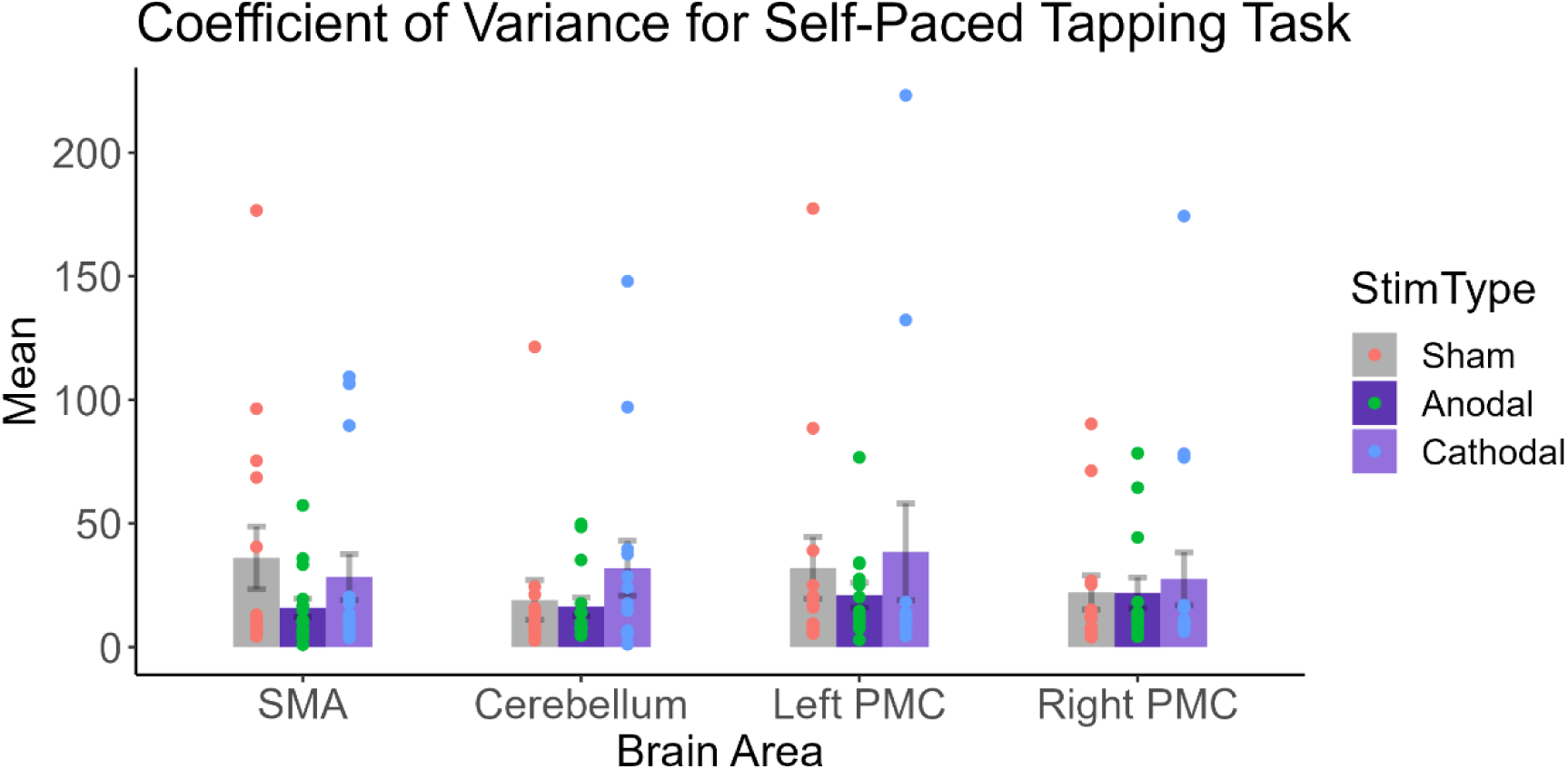
Self-paced tapping task. Mean Coefficient of Variance for the four groups (SMA, right cerebellum, left PMC and right PMC). Sham is in grey, anodal dark purple and cathodal in light purple. Individual coefficient of variance is shown by data points in orange for sham, green for anodal and blue for cathodal stimulations. No significant difference was seen between stimulation types and groups.

## Discussion

In this study, we investigated the causal role of four brain areas: SMA, right cerebellum, left PMC, and right PMC in beat perception. We examined how rhythm reproduction was affected by both anodal and cathodal stimulation compared to sham. Participants received sham, anodal, and cathodal stimulations in a counterbalanced order on different days. On each day, 20 trials each of strong-beat, weak-beat, and non-beat rhythms were reproduced. Participants also performed a self-paced tapping task while receiving stimulation. We hypothesized that modulating excitability in the SMA should influence the ability to reproduce strong-beat rhythms accurately, while altering the excitability of the cerebellum and PMC should influence the ability to reproduce weak and non-beat rhythms, or more non-beat rhythms when compared to the SMA stimulation. As predicted, participants had an overall better performance for strong-beat rhythms than for weak- and non-beat rhythms, and it was particularly better for high musical experience participants than low musical experience ones, replicating previous findings (Drake, 1993; Grahn & Brett, 2007). However, contrary to our main hypotheses, modulation of SMA, PMC, and cerebellum had no effect on reproduction accuracy for either beat-based or non-beat-based rhythms and had no impact on self-paced tapping variability. The analysis of averaged error also confirmed the null effects of tDCS on participant’s overall performance, with no decreased error following anodal stimulation and no increased error following cathodal stimulation. Bayesian statistical analysis revealed that beat strength and musical experience are the factors that most contributed to the improvement in rhythm reproduction performance, besides showing little evidence for the tDCS effects on the reproduction of rhythms.

From a behavioral standpoint, memory likely influenced participant’s performance as they had to remember each rhythmic sequence before reproducing the rhythms. Evidence supports the importance of working memory, which involves the use of short-term memory in an oriented task, for rhythm reproduction tasks (Saito & Ishio, 1998). Our task may have imposed a higher cognitive load, which is closely related to working memory and refers to the amount of information one can retain at a given time (Bannert, 2002). It might be the case where cognitive load was enough for participant’s recruiting other neural resources for the rhythm reproduction task so that the influence of SMA/PMC/cerebellum was less impactful in performance.

While anodal tDCS over the auditory cortex has been shown to affect memory for melodies (Schaal et al., 2021), the impact of tDCS over other brain regions related to beat perception such as the SMA, cerebellum and PMC, on memory for rhythmic sequences has not been demonstrated. No only that, previous research has demonstrated that moving to the beat improves overall listener’s timing perception (Manning & Schutz, 2013), and our data suggest that allowing participants to tap the rhythm in a rhythm reproduction paradigm is sufficient for improving overall performance, probably overriding any effect of stimulation that could arise when there is the need of any motor output.

### Motor brain regions and their roles in beat and time perception

#### Supplementary Motor Area

Here, although we did not find any effect of brain stimulation in the rhythm reproduction performance, beat strength played an important role for rhythm reproduction accuracy, and specifically for this group, participants with high musical experience had a better performance when reproducing the rhythms with a strong beat than participants with low musical experience. This might be because the SMA group had the largest number of participants with high musical experience (N = 14), when compared to participants with low musical experience (N = 8).

The absence of a causal effect within the SMA is surprising, particularly in light of the substantial body of research that has consistently highlighted the SMA’s involvement in functions such as rhythm perception, beat perception, and broader motor activities (Grahn & Brett, 2007, 2009; Leow et al., 2022). The SMA is involved in various motor activities, including sequencing actions, learning new motor abilities, and movement control in distracting situations (Nachev et al., 2008; Nguyen et al., 2014; Vollmann et al., 2013). Apart from that, the SMA’s networks, specifically the striato-thalamo-cortical loops, are essential for temporal predictions and the production and discrimination of time intervals (Macar et al., 1999, 2004). Thus, SMA helps anticipate the next beat in a sequence by sending direct signals to the dorsal striatum, which then creates representations of beat cycle intervals, thus closing this network loop through the activation of new SMA neural subpopulations via the thalamus (Cannon & Patel, 2021). Consequently, the SMA primarily supports beat-based timing sequences by planning the placement of the next beat in a sequence and maintaining the beat.

Studies where participants are required to synchronize their actions or responses with an external, predefined rhythmic cue (synchronization), and then are asked to continue the same rhythmic pattern without the external cue through an internal generation (continuation) support SMA’s importance for timing in a sequence; as SMA is important for the continuation phase, where individuals must rely on internalized timing information to maintain a rhythmic pattern (Halsband et al., 1993; Lewis et al., 2004; Rao, 1997). This aligns with the idea that the SMA is involved in generating the next beat in a sequence (Cannon & Patel, 2021). Neuroimaging studies have shown that the SMA exhibits higher activation during the continuation phase compared to the synchronization phase, indicating that the SMA is actively engaged in the internal generation of rhythmic patterns (Lewis et al., 2004; Rao, 1997). Moreover, patients with SMA lesions are impaired at continuation but not synchronization part of a task (Halsband et al., 1993; Rao, 1997). This suggests that the SMA is particularly crucial for tasks that require internal context-based timing processes (Goldberg, 1985). Considering these findings, it indeed becomes surprising that tDCS had no impact in your study, as it was expected to modulate the SMA’s activity during our reproduction task, that although different from the continuation tasks, maintain some similarities, such as the need to internally generate a beat. This discrepancy between expectations and outcomes highlights the complex nature of brain-behavior relationships and the need for further research using tDCS and other types of tasks such as the synchronization-continuation paradigm. In contrast to synchronization-continuation studies, our participants had to remember each sequence before reproducing the rhythms.

The main objective of our study was to extend the findings of a previous tDCS study on rhythm discrimination (Leow et al., 2022). Unlike in our data, Leow et al. (2022) found that SMA stimulation affected rhythm discrimination, with anodal stimulation improving performance and cathodal stimulation worsening it compared to sham stimulation. However, these results did not solely support SMA’s role in beat-based timing, as both strong and weak beat rhythms were equally affected by stimulation. It is possible that weak-beat rhythms were not irregular enough to affect the non-beat-based timing system, and the brain processed them using beat-based timing. To address this, we included non-beat rhythms to ensure that participants did not use beat-based timing in at least one condition. However, regardless of stimulation, we did not observe any performance differences among the different types of rhythms.

Considering the comparison to the rhythm discrimination results, the lack of differences in our rhythm reproduction task may stem from the fact that the SMA’s role in timing is more evident in perceptual tasks. Movement may recruit other supporting brain areas, masking the subtle temporal changes produced by SMA stimulation. When individuals engage in motor tasks or activities that involve the SMA, the complexity of the neural networks involved can make it challenging to isolate the specific effects of SMA stimulation on temporal processing, as supporting brain areas may compensate for any subtle temporal changes induced by SMA stimulation. While the SMA is implicated in timing and rhythm generation, the simultaneous involvement of other brain regions in motor control makes it challenging to pinpoint the exact impact of SMA stimulation on temporal changes. Alternatively, the SMA may not be responsive to the specific task of rhythm reproduction used in our study. Future research could incorporate non-beat rhythms into rhythm discrimination studies to test the hypothesis that stimulation of the SMA selectively affects beat-based timing in the context of less (weak-beat rhythms) and more (non-beat rhythms) complex rhythmic sequences.

#### Cerebellum

Contrary to our predictions, tDCS applied to the right cerebellum did not have any effect on the reproduction of non-beat rhythms or any other rhythms. This lack of observed benefits or costs suggests a null effect on the absolute timing system. Previous research on cerebellar stimulation has yielded conflicting results (Oldrati & Schutter, 2018; van Dun et al., 2016), potentially due to anatomical differences between the cerebellum and the cerebral cortex, because despite its small size, the cerebellum contains a majority of the brain’s neurons, organized differently from the cortex (Herculano-Houzel, 2009). Consequently, polarity differences (anodal versus cathodal) in tDCS effects are less predictable, making it challenging to anticipate the direction of behavioral changes (Oldrati & Schutter, 2018; Woods et al., 2016). In our study, we did not observe any effect of stimulation, regardless of whether it was anodal or cathodal.

Previous studies have demonstrated tDCS effects independent of the type of stimulation applied (anodal or cathodal) across various cognitive and motor tasks, including reaction time, working memory, motor learning, and motor memory (Ferrucci et al., 2008; Shah et al., 2013; Taubert et al., 2016). Null effects of cerebellar tDCS have also been widely reported in tasks involving associative learning, working memory, implicit learning, motor adaptation, and cognitive function (Beyer et al., 2017; Maldonado et al., 2019; Van Wessel et al., 2016; Verhage et al., 2017). These findings indicate that tDCS applied to the cerebellum does not consistently produce significant changes in performance across different tasks.

Although our study found a null result for cerebellar stimulation, it does not diminish the importance of the cerebellum in motor control, movement precision (Glickstein & Doron, 2008; Salman, 2002), and the absolute timing system (Nozaradan et al., 2017; Teki et al., 2011). It is possible that the specific rhythm reproduction task used in our study may not effectively engage the cerebellum. Future research should align cerebellar tDCS studies with neuroimaging techniques to confirm accurate targeting of the stimulation. Additionally, considering the entire cerebellum and exploring the effects of left cerebellar stimulation would be valuable. There is evidence of hemispheric specialization in the cerebellum, with some functions being more lateralized to the left or right hemisphere. For example, some studies suggest that the left cerebellum may be more involved in aspects of the singing activity, while the right cerebellum appears to be related to speech processing, and production of singing as well (Callan et al., 2006, 2007). By exploring both hemispheres separately, researchers can gain a more nuanced understanding of how tDCS may affect specific functions associated with each hemisphere during beat perception processing.

#### Premotor Cortex

Similarly to our cerebellar stimulation predictions, here we expected to see tDCS effects in weak-beat rhythms and non-beat rhythms more specifically when stimulating the premotor cortex, but no effect of anodal or cathodal stimulation was seen for either left or right premotor cortices. Previous research suggests that the PMC primarily plays a role in motor control and movement synchronization in response to external cues, rather than beat perception (Leow et al., 2022), and this might be the reason why we had no significant effect of stimulation over the reproduction of non-beat rhythms.

There is evidence linking the PMC to both beat-based and non-beat based, or absolute, timing, particularly in studies of rhythm synchronization (Chen et al., 2006, 2008a, 2008b, 2009). Different subregions of the PMC have been found to serve distinct roles in rhythm synchronization. For example, the ventral PMC is activated when participants listen to rhythms before synchronizing to them, while the dorsal PMC is responsive during synchronization, especially with more complex rhythms. The mid-PMC, along with the SMA and cerebellum, is activated when participants listen to rhythms without the intention of performing motor actions (Chen et al., 2008a). Additionally, the dorsal PMC has been shown to exhibit increased activation when the beat of a rhythm is made more salient (Chen et al., 2006). These findings highlight the involvement of the PMC in motor-auditory interactions during movement sequencing.

The null effects observed in our study are not entirely surprising considering the PMC’s general role in motor-time synchronization, and its activation in studies where no motor output, such as rhythm reproduction, is required. It is possible that our task failed to incorporate a crucial aspect of the PMC’s function, namely motor synchronization to an auditory stimulus or just passively listening to rhythms with no intention of tapping to the beat. In our study, participants did not have the opportunity to synchronize their movements to the rhythmic sequences but rather had to reproduce each rhythm from memory after hearing it multiple times. PMC’s causal role in beat perception through judgment tasks has been already shown to be null (Leow et al., 2022). Future research should investigate the effects of tDCS on the PMC in a rhythm synchronization paradigm to gain further insights into its role in temporal processing and motor coordination.

### Limitations

One limitation of our study and the field in general is the absence of a control task that has been previously shown to yield consistent results for SMA stimulation in a timing task. A control task that is consistently and predictably affected by stimulation has the benefit of adding a ground-truth to stimulation studies – they tell us whether (and in which subjects) tDCS was successfully applied. Here we tested whether self-paced tapping could be used as a control task but failed to demonstrate any effects of stimulation. While this could suggest that tDCS over the four brain areas studied here does not affect self-paced rhythm or motor output, it also means that we cannot confirm our predictions or validate the efficacy of stimulation in the sample. To address this, future research could explore different control tasks tailored to each specific brain area. If stimulation produces consistent effects in these control tasks but not in our rhythm reproduction task, it will support the conclusion that the four brain areas do not have a causal role in beat perception during rhythm reproduction.

One might blame our null results on the lack of categorization of participants according to musicianship, into musicians and non-musicians, instead of people with high and low musical experience. Future work could help to determine whether the lack of stimulation effects (or null effects) is due to task difficulty or the impact of stimulation itself incorporating a pre-task of rhythm reproduction to categorize participants as “strong beat perceivers” (higher accuracy) or “weak beat perceivers” (lower accuracy), and test for differences between groups.

Another important limitation is the use of a single-blind design where only participants were unaware of the type of stimulation received. Blinding tDCS is challenging due to the tingling and itching sensations often experienced by participants (O’Connell et al., 2012; Poreisz et al., 2007). Despite this limitation, our study was good in maintaining participant’s blindness towards the type of stimulation given the lower confidence ratings of the type of stimulation received.

Additionally, participants were not evenly distributed across the four groups based on their musical experience, and this may have influenced our results. The SMA group had more participants with high musical experience compared to other groups. Although participants were counterbalanced for the stimulated brain area, musical experience was not counterbalanced. Future studies should aim for an even distribution of participants with varying levels of musical experience.

Addressing the limitations of control tasks, task complexity, blinding, and participant characteristics will enhance the validity and generalizability of future research investigating the effects of tDCS on beat perception.

## General Conclusions

Previous research has demonstrated that moving to the beat improves overall listener’s timing perception; movement is not only important for ‘time acuity’, or the precision in perceiving short time intervals but also to the ‘timekeeping’, which relates to the broader task of tracking and maintaining time over longer durations, allowing more accurate detection of timing deviations (Manning & Schutz, 2013). As demonstrated by the differences in reproduction according to the beat strength, our data suggest that allowing participants to tap the rhythm in a rhythm reproduction paradigm is sufficient for improving overall performance, probably overriding any effect of stimulation that could arise.

A null result like the one we obtained in our rhythm reproduction task cannot lead us to conclude that the SMA is not necessary for beat perception. Beat perception can be measured by different tasks and functions and taken together with the results of tDCS in the SMA during rhythm discrimination – the effects of stimulation may be too weak to be observed during reproduction. Similarly, effects of stimulation on rhythm discrimination were seen for the cerebellum (Leow et al., 2022), but not during reproduction here. The different roles of motor brain regions in rhythm perception and production therefore may be easier to observe with tDCS when more sensory, rather than sensorimotor, tasks are used.

